# During environmental change, cooperation can promote rescue or lead to evolutionary suicide

**DOI:** 10.1101/784553

**Authors:** Gil J. B. Henriques, Matthew M. Osmond

**Affiliations:** Department of Zoology & Biodiversity Research Centre, University of British Columbia, Canada; Center for Population Biology, University of California Davis, USA

**Keywords:** cooperation, public goods, moving optimum, environmental change, evolutionary rescue, evolutionary suicide

## Abstract

The adaptation of populations to changing conditions may be affected by interactions between individuals. For example, cooperative interactions may allow populations to maintain high densities, and thus keep track of moving environmental optima. At the same time, changes in population density alter the marginal benefits of cooperative investments, creating a feedback loop between population dynamics and the evolution of cooperation. Here we model how the evolution of cooperation is affected by, and in turn affects, adaptation to a changing environment. We hypothesize that changes in the environment lower population size and thus promote the evolution of cooperation, and that this in turn helps the population keep up with the moving optimum. However, we find that the evolution of cooperation can have qualitatively different effects, depending on which fitness component is reduced by the costs of cooperation. If the costs decrease fecundity, cooperation indeed speeds adaptation by increasing population density; if, in contrast, the costs decrease viability, cooperation may in fact slow adaptation by lowering the effective population size, leading to evolutionary suicide. Thus, we show that cooperation can either promote or—counter-intuitively—hinder adaptation to a changing environment.

## 1. Introduction

Throughout evolutionary history, populations have experienced constantly changing environmental conditions. Examples of large-scale environmental changes with profound evolutionary legacies include the rise of atmospheric oxygen during the Proterozoic (Lyons and Reinhard, 2014); continental movements like the emergence of the Isthmus of Panama during the Pliocene (Stehli and Webb, 1985); and climatic alterations such as the Pleistocene glaciations (Hewitt, 2004). In recent times, natural populations are increasingly facing anthropogenic directional environmental shifts, ranging from habitat degradation and climate change (reviewed in Gienapp et al. 2008; Hoffmann and Sgrò 2011; Sih et al. 2011) to exposure to increasing concentrations of antibiotics (Klein et al., 2018).

Because populations are often locally adapted (Hereford, 2009), environmental changes may lead to decreased fitness and to a decline in population size. Ultimately, this may lead to extinction, as the population’s phenotype is unable to track the shifting environmental optimum. This fate may be prevented when adaptation counters population decline and prevents extinction, a process called evolutionary rescue (Bell, 2017; Gomulkiewicz and Holt, 1995). The effects of environmental change (and ultimately the possibility of evolutionary rescue) will, in general, depend on within–and between-species biotic interactions (Collins, 2011; Harmon et al., 2009; Lavergne et al., 2010; Lawrence et al., 2012; Tylianakis et al., 2008). Although most theoretical models of adaptation to changing environments ignore ecological interactions—reviews of theory by Alexander et al. (2014) and Bell (2017) barely discuss the subject, only very briefly mentioning competition—there is a growing body of modeling work on the effects on evolutionary rescue of ecological interactions, both between and within species, such as competition (e.g., Johansson 2007; Osmond and De Mazancourt 2012) and predation (e.g., Jones 2008; Mellard et al. 2015; Osmond et al. 2017).

Here we focus on how adaptation to a changing environment is affected by a different type of interaction, cooperation. Because high levels of cooperation can increase population abundance (Chuang et al., 2009; Tekwa et al., 2017)—and larger populations are both less prone to extinction from demographic stochasticity and can adapt faster due to a larger mutational input and less drift—we predict that cooperation can reduce the chance that a population will go extinct due to environmental change.

Following a well-established game-theoretical tradition (reviewed in Archetti and Scheuring 2012), we will model cooperative interactions as public goods games: events in which individuals pay a cost to contribute toward a public good. The benefit generated by this public good is then equally enjoyed by all interacting partners. Public goods interactions can describe many instances of cooperative behavior in nature (Levin, 2014). For example, many microbes secrete extracellular molecules which can be utilized by non-producers (Tarnita, 2017; West et al., 2006); these include, for example, adhesive polymers in *Pseudomonas* (Rainey and Rainey, 2003), invertase in yeast (Gore et al., 2009), and indole (a signaling molecule providing antibiotic resistance) in *Escherichia coli* (Lee et al., 2010). Similarly, in many costly or risky behaviors such as cooperative breeding (Rabenold, 1984), cooperative hunting (Bednarz, 1972; Packer et al., 1990; Yip et al., 2008), alarm calls against predators (Clutton-Brock et al., 1999), or the formation of fruiting bodies in social amoebae (Strassmann et al., 2000), benefits are equally distributed to all participants. Even cancer cells share costly diffusible products, such as growth factors (Archetti et al., 2015; Axelrod et al., 2006). Thus, public goods interactions are a useful general framework to model cooperation, applicable to a wide range of natural scenarios.

The public goods framework also permits a seamless articulation between evolutionary game dynamics, on the one hand, and ecological or demographic dynamics (such as changes in population size, as might be caused by environmental shifts), on the other. Models that make this connection explicit (called ‘ecological public goods games’, Gokhale and Hauert 2016; Hauert et al. 2006, 2008; Parvinen 2010; Wakano et al. 2009) are based on the feedback between population size and the fitness of cooperators. In public goods interactions, the marginal benefits of cooperation decrease with higher group sizes, meaning that cooperation is favored only when groups are small. This pattern matches empirical observations; for example, the amount of time that meerkats spend on guard decreases with group size (Clutton-Brock et al., 1999), and *Pseudomonas* ‘cheaters’ that do not produce siderophores perform better at high cell densities (Ross-Gillespie et al., 2009). Therefore, if low population densities correspond to small interaction group sizes, they allow cooperation to gain a foothold. As cooperation increases, so does the population mean fitness, and, consequently, the population density. This feedback loop between ecology and evolution can lead to the maintenance of stable, intermediate frequencies of cooperators (Hauert et al., 2006, 2008). Laboratory experiments in sucrose-growing yeast (Chen et al., 2014; Harrington and Sanchez, 2014; Sanchez and Gore, 2013) confirm the empirical relevance of these feedbacks between population density and the evolution of cooperation. Populations whose growth is mediated by cooperative production of invertase approach intermediate densities and frequencies of cooperators, consistent with ecological public goods dynamics (Sanchez and Gore, 2013).

Cooperation in public goods games was initially described (Hamburger, 1973) in the context of bimorphic populations where individuals are either cooperators (who produce a fixed amount of public good) or defectors (non-producers). For example, whereas wild-type yeast cells produce invertase, mutant ‘cheaters’ do not (Gore et al., 2009). This is also the approach used in most models of ecological public goods games (Hauert et al., 2006, 2008; Wakano et al., 2009). However, it can also make sense to imagine cooperation as a quantitative trait. For instance, meerkat females can provide more or less assistance with babysitting and pup feeding (Clutton-Brock et al., 2001). In continuous public goods games (Doebeli et al., 2004; Killingback and Doebeli, 2002) individuals are described by a continuous trait value that quantifies how much they invest into the public good. Parvinen (2010) showed that, if cooperation is modelled as a continuous trait, ecological public goods games can exhibit evolutionary branching—that is, the population can evolve into a highly cooperating and a non-cooperating strain. In other words, the continuous public goods framework can also (depending on the parameters of the model) give rise to discrete cooperators and defectors, a process called ‘tragedy of the commune’ by Doebeli et al. (2004), who first described it in the context of the continuous snowdrift game. In our model, we will use the continuous public goods game. As we will see, depending on the shape of the function that describes the costs of cooperation, we may observe a tragedy of the commune; if this happens, the model will become identical to a discrete public goods game. Thus, the same framework will allow us to investigate the interaction between cooperation and environmental change, both in those cases where cooperation is continuous and in those where it is discrete.

Under directional environmental change, species keep track pf the moving environmental optimum at a certain phenotypic distance or ‘lag’ (Bürger and Lynch, 1995; Lynch and Lande, 1993). The size of the lag will depend on the speed of environmental change, as well as the population’s ability to adapt (Bürger and Lynch, 1995; Lynch and Lande, 1993). The lag reaches an equilibrium when the rate of environmental change equals the rate of adaptation, which, for mutation-limited evolution, is proportional to population size and mutation rate (Dieckmann and Law, 1996). The larger the lag, the lower the size of the population. If the lag becomes too large, the population will get trapped in a vicious cycle: it will grow so small that it cannot adapt fast enough to keep up with the environmental optimum. The result is an extinction vortex (Brook et al., 2008; Fagan and Holmes, 2006; Gilpin and Soulé, 1986): an amplifying feedback that will ultimately lead the population to extinction (Johansson, 2007); in the quantitative genetics literature, a similar cycle exists with genetic variance taking the place of population size (Bürger and Lynch, 1995; Osmond and Klausmeier, 2017). However, a population engaged in ecological public goods games could potentially break free from this vicious cycle, because the decline in population density would favor cooperation. This matches experimental evidence that, as environments deteriorate, public goods producers increase in frequency (Chen et al., 2014). Thus, our hypothesis is that environmental change favors the evolution of cooperation, and that this process may rescue a population that would otherwise have been unable to keep track with the optimum.

## 2. Methods and results

As a modeling framework, we will use adaptive dynamics (Dieckmann and Law, 1996; Geritz et al., 1998; Metz et al., 1996) to follow the change, over time, of a population whose fitness depends on the match between a given functional quantitative trait *y* ∈ ℝ and the environment (an approach similar to Johansson 2007). Apart from their functional trait, individuals are also characterized by a cooperation trait 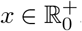, describing the amount of investment into public good production. These traits are completely genetically determined and individuals are assumed to reproduce asexually. It would be interesting to extend the model to include environmental noise and plasticity, as well as alternative models with diploidy and sexual reproduction.

The population, of size *n*, is assumed to be quasimonomorphic, consisting mostly of resident individuals characterized by a vector **z** = (*x*, *y*). A very small fraction of individuals carry small-effect mutations in these traits. Because mutants are rare, we neglect their contribution to the environment experienced by any focal individual. Trait values of focal individuals are distinguished with a prime: **z**′ = (*x*′, *y*′).

### A. Fitness components

Generations are nonoverlapping, i.e., individuals are semelparous (they reproduce only once) and reproduce synchronously by producing clutches, litters, or broods (as may be the case with many bee and wasp species, whose life-cycles are annual). A focal individual with phenotype **z**′, in an environment that is set by the resident with phenotype **z**, may survive to reproductive age with a probability equal to their viability, *V* (**z**′|**z**) (otherwise, they die before reproducing, but after a large number of interactions). Should they survive, they will give birth to an expected number of offspring equal to their fecundity, *F*(**z**′|**z**), and die after giving birth. Thus, the contribution of a focal individual to the next generation is determined by two random variables: survival to reproductive age (*X*)—which is Bernoulli-distributed with expectation *V* (**z**′|**z**)—and the number of offspring after reproduction (*Y*)—which is assumed to be Poisson-distributed with expectation *F* (**z**′|**z**).

The contribution of a focal individual to the next generation is drawn from the product of the two random variables, *XY*, whose expectation is the individual’s Wrightian fitness, *W*(**z**′|**z**) ≡ E[*XY*] = *V* (**z**′|**z**) · *F*(**z**′|**z**). The variance of the same distribution is given by 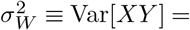 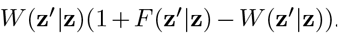. At ecological equilibrium (when *W*(**z**|**z**) = 1), this quantity, which we will call the variance in reproductive success, is equal to 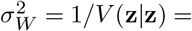 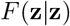. This variance measures the amount of stochasticity in reproductive success, given a particular trait value, and thus affects the relative strengths of selection and drift (i.e., the effective population size).

During their lifetime, individuals engage in many interactions, from which they collect costs and benefits. An individual’s baseline (maximum) fecundity is equal to the lifetime average benefits of cooperation collected by that individual (*B̅*(*x*′|*x*, *n*), section A.1); the maximum viability is one. Other life-cycle factors—viz., the costs of cooperation (section A.1), the mismatch between the individual’s phenotype and the environment (section A.2), and density regulation (section A.3)—will decrease these baseline values. We assume no deaths occur until individuals have participated in many interactions, such that fitness only depends on the current population size *n*, the average benefits of cooperation, and the three life-cycle factors enumerated above.

#### A.1. Public goods games

Each interaction begins with a focal individual assembling an interaction group. Every individual in the population has the same probability *p* of joining the interaction group (i.e., there is no spatial structure or heritable variation in interaction tendency). For example, there may be a probability *p* of being inside the diffusion range of a given growth factor. Thus, the probability Pr(*g*|*n*) that the interaction group size will equal *g* is given by:

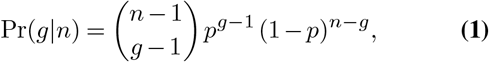

which, for integer arguments, is the binomial distribution. Generalizing the binomial coefficient, 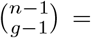 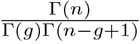, Eq. 1 applies for noninteger arguments as well.

Eq. 1 mechanistically connects population dynamics to public goods dynamics. Variations in population size, *n*, will affect the group size, which in turn (see below) changes the incentives for cooperation. This feedback loop promotes the maintenance of cooperation at intermediate levels, and it makes our model an example of an ecological public goods game (Gokhale and Hauert, 2016; Hauert et al., 2006, 2008; Parvinen, 2010; Wakano et al., 2009). Whereas previous models implemented eco-evolutionary feedbacks by limiting reproductive opportunities, thus implicitly incorporating space (Gokhale and Hauert, 2016; Hauert et al., 2006, 2008; Parvinen, 2010; Wakano et al., 2009), Eq. 1 accomplishes the same qualitative effect via a different mechanism (by assigning each individual a probability of joining interaction groups).

In each interaction, every individual contributes to a common pool of public good. The total quantity is multiplied by a multiplication factor *r*, generating a benefit that is equally distributed among the members of the group. Therefore, for each interaction, the benefit to a focal individual with trait value *x*′ in a group of *g* − 1 resident individuals is:

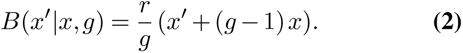

Over the course of a lifetime, the focal individual will have participated in many interactions, collecting an average benefit *B̅*:

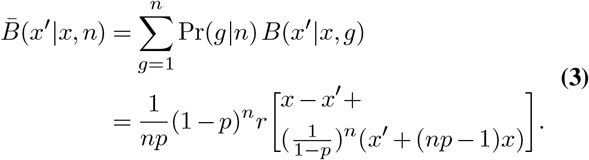

Any nonzero amount of investment into producing public goods carries a fitness cost. The focal actor with phenotype *x*′ will incur a cost *C*(*x*′); the relevant fitness component (viability or fecundity) is multiplied by the complement of the cost, (1 − *C*(*x*′)). Although the cost increases with the amount of investment, we assume that the rate of this increase gets smaller as the investment level grows: lim_*x*′→∞_*C*(*x*′) = 1. This assumption is realistic in the case of economies of scale. We will use the following functional form:

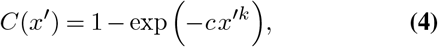

where higher values of *c* and *k* imply faster and more concave increases of cost with investment, respectively.

Since the cost depends only on the focal individual’s phenotype, the average cost is simply equal to the cost for one interaction.

#### A.2. Environmental mismatch

The state of the environment, *θ*, is measured in the same scale as the functional trait, *y*.

We assume fitness is maximized when *y* = *θ*, and explore the case where *θ* changes at a constant velocity, *v*. With a moving environment, there will be a mismatch between the trait value *y* and the optimum *θ*, which decreases fitness. In this case, we say that the population exhibits a lag, 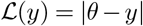, and the relevant fitness component is multiplied by the Gaussian function *M*(*y*′):

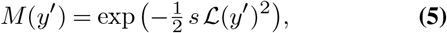

where *s* represents the strength of stabilizing selection on *y*.

#### A.3. Density regulation

We implement density regulation similarly to Beverton and Holt (1957). The relevant fitness component is multiplied by a decreasing function *D*(*n*) of population size:

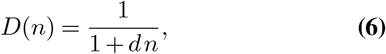

where *d* determines the rate at which fitness decreases with population size.

### B. Resident dynamics

In order to study the evolution of a population adapting to a moving optimum, we will apply a time-scale separation. Following standard adaptive dynamics methodology, we assume that evolution is much slower than ecological population dynamics (Dieckmann and Law, 1996; Geritz et al., 1998; Metz et al., 1996). This will be the case, for example, if evolution is limited by rare small-effect mutations, rather than occurring by standing genetic variation. This time-scale separation allows the population to always reach ecological equilibrium (i.e., a monomorphic population at carrying capacity) before the introduction of new advantageous phenotypes. We also assume that environmental change occurs on an evolutionary timescale (as in Johansson 2007), i.e., during the time it takes for a population to reach ecological equilibrium the environmental state changes only by a negligible amount. Note that if the environment were to change much faster than the evolutionary timescale, then evolutionary rescue would be impossible; if it were to change much slower, then it would be trivial.

For a given population size *n*, we can calculate the inter-generation change in the expected population size: Δ*n* = *n W*(**z**|**z**) − *n*, where *W*(**z**|**z**) = *B̅*(*x*|*x*, *n*)(1 − *C*(*x*))*D*(*n*)*M*(*y*). When the population size is constant, Δ*n* = 0, the population is at ecological equilibrium and will have size *n̂*(**z**)(Fig. 1A):

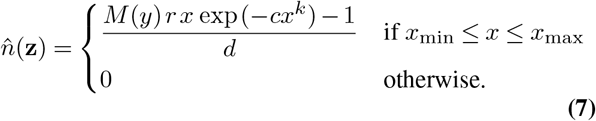

**Fig. 1.**
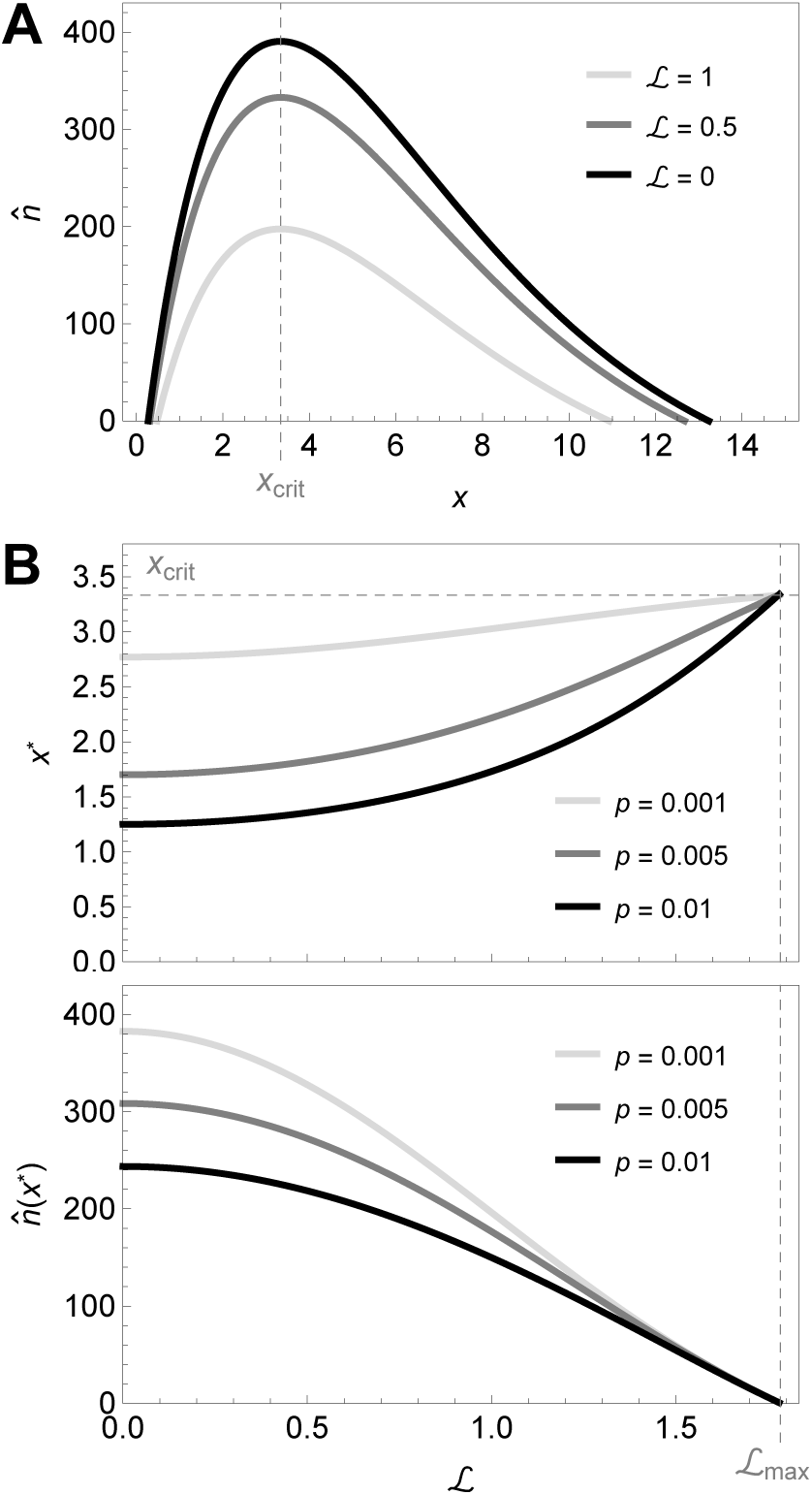
The population size depends on distance to the optimum and on the amount of cooperative investment. **A:** The population size at ecological equilibrium (*n̂*) increases (up to a point) with cooperation *x* and decreases with distance 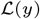 to the environmental optimum (Eq. 7). At very low values of cooperative investment (Eq. 8) or very large lags (Eq. 10), the population goes extinct. The population size is maximized when *x* = *x*_crit_ (Eq. 9). **B**, *Top:* the equilibrium value of cooperation with a constant lag (*x*^⋆^, calculated from Eq. 11) increases for higher lags 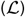 and smaller interaction group sizes (smaller *p*). *Bottom*: corresponding population size at equilibrium (*n̂*(*x*^⋆^)). Parameters: *d* = 0.01, *s* = 1, *c* = 0.3, *r* = 4, *k* = 1, *p* = 0.01 (except when otherwise stated in the figure).

As expected, the population size is higher when public goods are cheaper to produce (low *c*) or provide higher benefits (high *r*), as well as when density regulation is less intense (low *d*). Furthermore, even in a constant environment, a minimum amount of cooperation is necessary to sustain the population and avoid extinction (Fig. 1A):

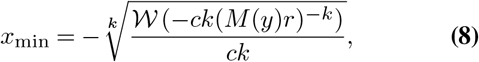

where 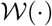 denotes the Lambert *W*-function, also known as the product logarithm (Corless et al., 1996; Lehtonen, 2016). As individuals invest more in cooperation, the population size increases (Fig. 1A), up to a critical value, *x*_crit_:

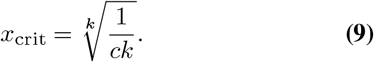

Beyond this point, the costs of cooperation are so high that further investment leads to smaller populations. Therefore, perhaps surprisingly, there is a maximum amount of cooperation, *x*_max_ beyond which the population cannot persist (Fig. 1A).

The equilibrium population size also depends on the lag between the population’s trait and the environmental optimum (Fig. 1A). Substituting Eq. 5 into Eq. 7 and solving *n̂*(**z**) = 0 for 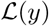, we can calculate the maximum value of lag above which the population goes extinct:

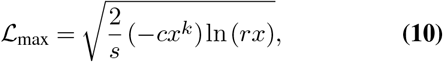

### C. Evolution of cooperation for a constant lag

We now focus on the fate of a rare mutant with trait value *x*′, and we will consider the case of a constant lag, e.g., 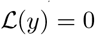, in a constant environment (*v* = 0). Because we are concerned with adaptation in *x* and the lag is fixed, we will drop the dependence on *y* for the remainder of this section. The mutant’s fitness is then given by *W*(*x*′|*x*) = *B̅*(*x*′|*x*, *n̂*(*x*))(1 − *C*(*x*′))*D*(*n̂*(*x*))*M*.

The direction and strength of selection on cooperation can be measured by the selection gradient of investment, which is given by

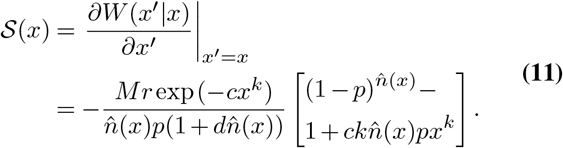

The evolutionary dynamics will be at equilibrium when there is no directional selection on cooperation. At this point, the selection gradient vanishes, *S*(*x*^⋆^) = 0, and *x*^⋆^ is said to be an evolutionarily singular point (Geritz et al., 1998). Although an explicit analytical solution is impossible to calculate, we can solve for *x*^⋆^ numerically. As expected, the equilibrium level of investment *x*^⋆^ is higher if the distance to the optimum is higher and if interaction groups are small (small *p*), Fig. 1B.

When *x* = *x*_crit_, the term in the square brackets in Eq. 11 becomes equal to 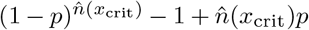, which is guaranteed to be positive, so that *S*(*x*_crit_) < 0. Therefore, in a monomorphic population, the singular point *x*^⋆^ will never take values higher than *x*_crit_ (Eq. 9). Intuitively, for any given lag, increasing cooperation beyond *x*_crit_ would lead to decreases in population size.

#### C.1. Convergence to the branching point and evolutionary stability

The singular point will be an attractor of the evolutionary dynamics if, starting from a strategy in the neighbourhood of the singular point, the population converges toward *x*^⋆^, i.e., 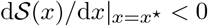 (Geritz et al., 1998). This can be visualized by drawing a pairwise invasibility plot (PIP), which indicates the mutant strategies *x*′ that can invade any given resident strategy *x* (Geritz et al., 1998; van Tienderen and de Jong, 1986). As the PIPs in Fig. 2 illustrate, the singular point in our model can be an attractor of the evolutionary dynamics, since mutant strategies can invade whenever they are closer to the singular point than the resident strategy. Thus, a monomorphic population will converge toward an intermediate amount *x*^⋆^ of cooperative investment.

**Fig. 2.**
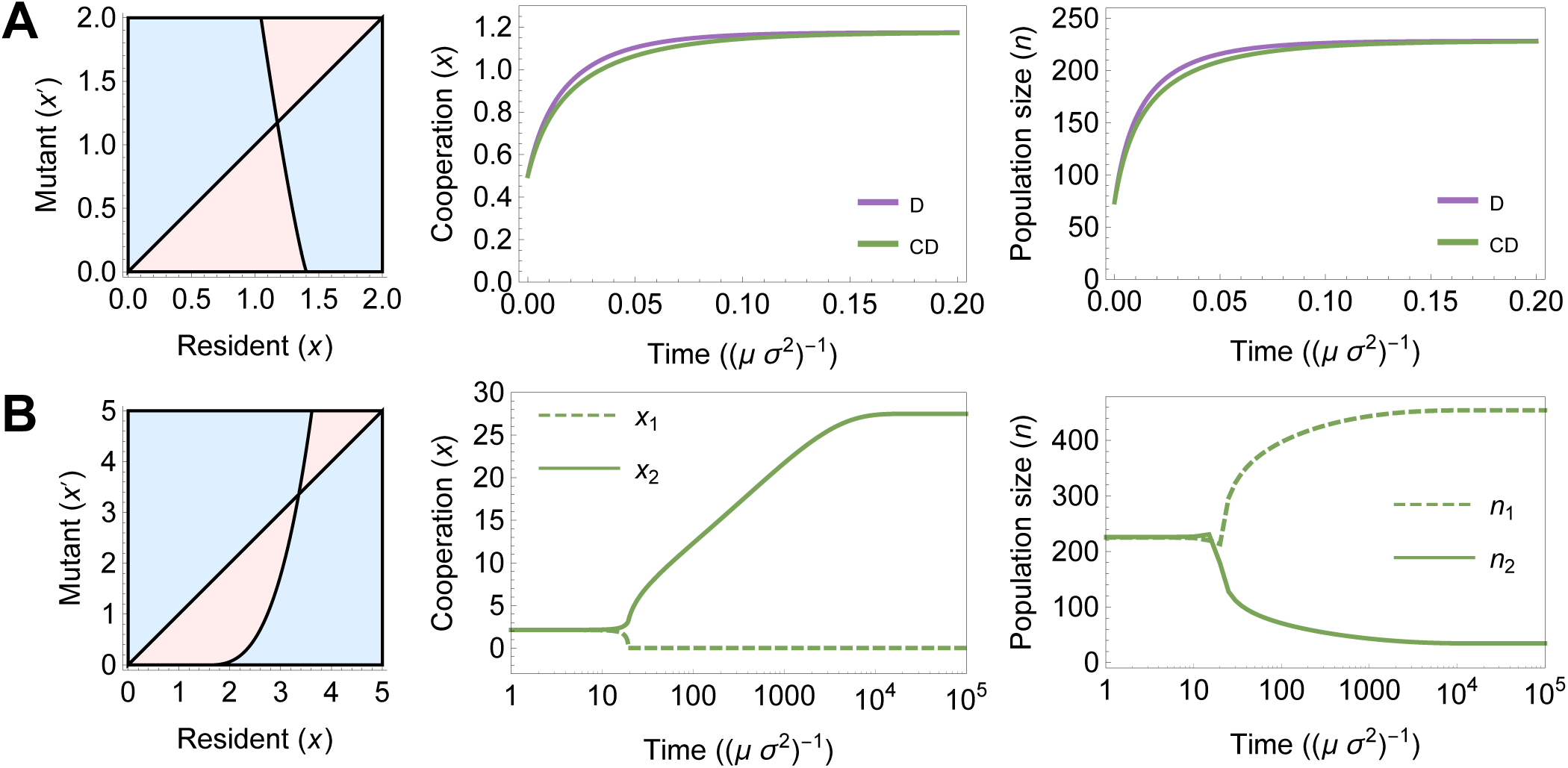
Monomorphic evolutionary dynamics for a constant lag when the singular point *x*^⋆^ is a stable point (**A**) and when it is a branching point (**B**). The leftmost panels are pairwise invasibility plots, where blue [red] areas correspond to mutants with higher [lower] fitness than the r esident. The point at which the black lines cross is the singular point; this point cannot be invaded by any mutants in **A**, but it is invasible on both sides in **B**. The other panels show the monomorphic evolutionary dynamics of cooperation (*x*, middle panels) and corresponding population size (*n*, rightmost panels). In **A**, a population starting away from the singular point evolves toward toward *x*^⋆^ and remains at equilibrium. Purple and green curves correspond, respectively, to the life-cycles D and CD. In **B**, only one life-cycle (CD) is shown; a population starting at singular point branches into a defector (*x*_1_) and a cooperator (*x*_2_) strain. Parameters: *d* = 0.01, *r* = 4, *c* = 0.3, *M* = 1. For **A**: *k* = 1.1; for **B**: *k* = 0.3.

Once the population reaches equilibrium, there are two possible outcomes. If the singular strategy is a fitness maximum, then evolution will come to a halt, and cooperation will remain constant at *x*^⋆^ (evolutionarily stable strategy). Otherwise (if the singular strategy is a fitness minimum), then the monomorphic population can simultaneously be invaded by mutants on either side of *x*^⋆^. This leads the population to split into two distinct and diverging phenotypes—a process called evolutionary branching (Geritz et al., 1998). Depending on the model parameters, *x*^⋆^ can either be evolutionarily stable (Fig. 2A) or unstable (Fig. 2B). For individual-based simulations illustrating both possibilities, see Figs. S1A (evolutionarily stable) and S1B (evolutionary branching) in the Supplemental Information.

The singular strategy will be a fitness minimum if:

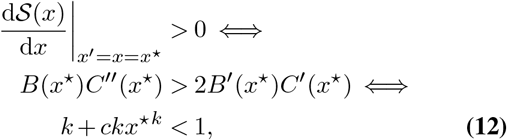

that is, the singular strategy will be a branching point only when the cost is sufficiently weak (small *c*) and convex (small *k*), and the equilibrium level of cooperation is low.

If the singularity is a branching point, the population branches into a defector strain and a cooperator strain (a process called tragedy of the commune, see Doebeli et al. 2004), with trait values *x*_1_ and *x*_2_ respectively (Fig. 2B). At evolutionary equilibrium, 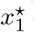 and 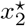 do not depend on the lag (as shown in section S2.1 of the Supplemental Information), and therefore the model becomes identical to a discrete public goods game, even in a changing environment. Hence, the same model allows us to investigate how the evolution of cooperation affects adaptation to changing environments in continuous as well as in discrete public goods games.

In this manuscript, we will focus in the case where the singular strategy is evolutionarily stable (Fig. 2A). Individual-based simulations suggest that, although branching is possible (Fig. S1B, Supplemental Information), the parameter range under which this process actually does occur, resulting in a stable dimorphic equilibrium, is relatively narrow when compared to the analytical predictions above. In particular the population size of the cooperator is often so small that stochastic extinction of that branch quickly occurs. Nonetheless, we present some remarks regarding the post-branching dynamics in section S2 (Supplemental Information).

#### C.2. Monomorphic evolutionary dynamics for a constant lag

The selection gradient can also be used to determine the rate of evolution. Using a traditional adaptive dynamics approach (Dieckmann and Law, 1996; Geritz et al., 1998; Metz et al., 1996), adapted for a life-cycle with non-overlapping generations (section S1 in the Supplemental Information), we obtain a differential equation for the time-dynamics of cooperative investment (*ẋ*):

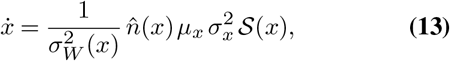

where *μ*_*x*_ and 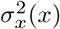 refer, respectively, to the probability of mutation at birth and to the variance of the mutational distribution for *x*, and 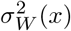 refers to the variance in the resident’s reproductive success (section A). Eq. 13 is analogous to the canonical equation of adaptive dynamics (Dieckmann and Law, 1996). In contrast to the canonical equation, the rate of adaptation in Eq. 13 is inversely proportional to the variance in reproductive success, 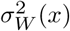. Intuitively, given the same expected number of offspring, variance in number of offspring increases the chance that an advantageous mutant is lost through genetic drift; therefore, variance in reproductive success slows adaptation down (Supplemental Information, see also Durinx et al. 2008).

Substituting Eqs. 7 and 11 into Eq. 13 we are able to determine the time-dynamics of cooperation. At this point, however, we need to be explicit about the details of the life-cycle of the species we are studying. Because the rate of adaptation depends on the variance in reproductive success, 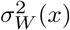, the evolutionary trajectory will depend on how the different life-cycle factors—environmental mismatch (Eq. 5), cost of public good production (Eq. 4), and density regulation (Eq. 6)—are partitioned among the two fitness components (viability and fecundity, see section A). Each of the the three life-cycle factors in our model can affect either viability or fecundity, for a total of eight possible life-cycles (Table 1). For example, the mismatch between the environmental state and the functional trait may conceivably decrease either viability or fecundity, depending on the species. The choice of life cycle will turn out to qualitatively affect the outcome of adaptation to changing environments, via its effect on variance in reproductive success 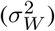.

**Table 1.**
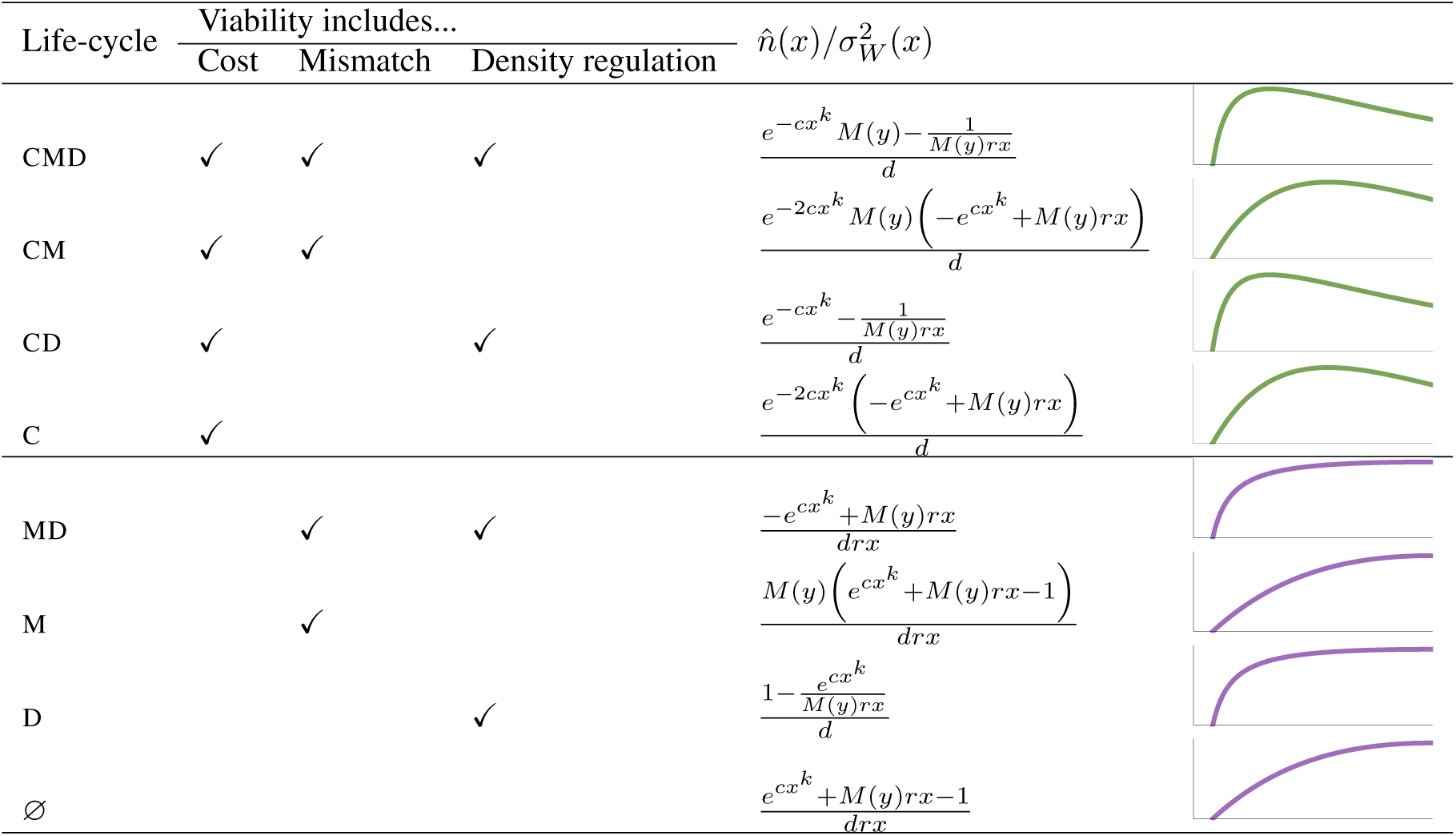
Possible life-cycles in the model. Life-cycles are named after the initial letter of the model elements (<u>c</u>ost of cooperation, environmental <u>m</u>ismatch, or <u>d</u>ensity regulation) that decrease viability. Elements that do not decrease viability decrease fecundity instead. Abscissae indicate cooperation level (0 < *x* < *x*_*crit*_, Eq. 9), and ordinates indicate 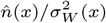. Parameters: *d* = 0.01, *r* = 4, *c* = 0.3, *p* = 0.01, *M* (*y*) = 1, *k* = 1.

| Life-cycle | Viability includes... | | | $\hat{n}(x)/\sigma_W^2(x)$ | |
| --- | --- | --- | --- | --- | --- |
|  | Cost | Mismatch | Density regulation |  |  |
| CMD | ✓ | ✓ | ✓ | $\frac{e^{-cx^k} M(y) - \frac{1}{M(y)rx}}{d}$ | |
| CM | ✓ | ✓ | | $\frac{e^{-2cx^k} M(y) (-e^{cx^k} + M(y)rx)}{d}$ | |
| CD | ✓ | | ✓ | $\frac{e^{-cx^k} - \frac{1}{M(y)rx}}{d}$ | |
| C | ✓ | | | $\frac{e^{-2cx^k} (-e^{cx^k} + M(y)rx)}{d}$ | |
| MD | | ✓ | ✓ | $\frac{-e^{cx^k} + M(y)rx}{drx}$ | |
| M | | ✓ | | $\frac{M(y) (e^{cx^k} + M(y)rx - 1)}{drx}$ | |
| D | | | ✓ | $\frac{1 - \frac{e^{cx^k}}{M(y)rx}}{d}$ | |
| ∅ | | | | $\frac{e^{cx^k} + M(y)rx - 1}{drx}$ | |

Two qualitatively different behaviors will be observed, depending on whether the costs of public good production decrease viability (green curves in Table 1) or fecundity (purple curves). Throughout the text, we will illustrate the effect of life-cycle choice by contrasting two example life-cycles (which we call CD and D, see Table 1). In life-cycle CD, the costs of public good production decrease viability, whereas in life-cycle D, they decrease fecundity. As we will see, which fitness component is affect by density regulation and environmental mismatch has no qualitative effect. In both example life-cycles, density regulation decreases viability, whereas environmental mismatch decreases fecundity (Table 1).

The viability of a resident individual with life-cycle D is *V*_D_(*x*|*x*) = *D*(*n̂*(*x*)) = exp(*cx*^*k*^)/(*Mrx*), and that of a resident individual with life-cycle CD is *V*_CD_(*x*|*x*) = (1 − *C*(*x*)) · *D*(*n̂*(*x*)) = 1/(*Mrx*). As evolution drives the population toward higher levels of cooperative investment, the costs of cooperation increase, so that CD-species adapt slower and slower when contrasted with D-species (Fig. 2A). As we will see later on, this will lead to qualitative differences in the capacity for evolutionary rescue, but in the case of adaptation toward a stable singular point with a constant lag the long-term outcome of evolution is the same (Fig. 2A).

### D. Evolution of cooperation during environmental change

In section C, we considered the dynamics of cooperation when the level of cooperative investment is evolving and the functional trait is at some constant distance to the environmental optimum. In the long term, populations that track moving environments will indeed always evolve to exhibit a constant lag (or, alternatively, they will fail to track the optimum and go extinct). However, the dynamics from section C do not take into account the feedback between cooperation and lag distance. We will now track how populations evolve while the environment changes at some velocity *v*.

We imagine that populations are initially well-matched to the environment (and at evolutionary equilibrium, see section C). Recall that, if the evolutionary singular point *x*^⋆^ is an evolutionary branching point (Eq. 12), then the initially well-adapted population will be participating in a discrete ecological public goods game (i.e., it will consist of two distinct strains: cooperators and defectors, whose frequencies may fluctuate as the environment changes; for more details see section S2.1 in the Supplemental Information). Otherwise, it will be engaged in a continuous ecological public goods game (i.e., it will be a monomorphic population characterized by a quantitative trait, describing the amount of cooperative investment). Here, we will consider the latter scenario; for some remarks on the prior case see section S2.2 in the Supplemental Information.

We again focus on the fate of rare mutants, with trait values **z**′ = (*x*′, *y*′). A mutant’s fitness is equal to *W*(**z**′|**z**) = *B̅*(*x*′|**z**) (1 − *C*(*x*′))*D*(**z**)*M*(*y*′). The formula for the selection gradient of cooperation is given by Eq. 11 (with the added dependence of *M* on *y*′). As for the functional trait, its selection gradient is given by:

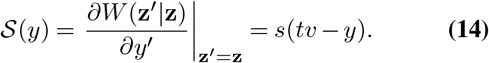

In contrast to section C.2, there is now no endpoint to evolution because, if the population manages to change fast enough to avoid extinction, it will approach a state in which *y* is constantly evolving at the same velocity as the environment (*ẏ* = *v*). Nonetheless, there may still be a ‘dynamical’ equilibrium where the lag stabilizes at some constant value. To find the dynamical equilibrium, we are ultimately interested in finding the values of *x* and 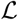 for which *ẋ* = 0 and 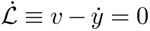. As there is no explicit analytic solution for Eq. 11, we will solve for *x* and 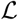 numerically. Then, we will further clarify our results by analytically determining the maximum velocity of environmental change that the population is able to track without going extinct.

The differential equation for the dynamics of the two traits **z** = (*x*, *y*), with selection gradients **S**(**z**) = (𝓢(*x*), 𝓢(*y*)), can be calculated similarly to Eq. 13:

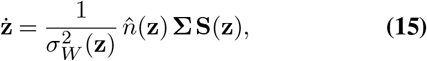

where 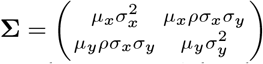 is the variance-covariance matrix for the mutational distribution. The coefficient *ρ* describes the correlation between mutations in the two traits (and will henceforth be set to zero); *μ*_*x*_ and *μ*_*y*_ are the probabilities, at birth, of a mutation in trait *x* or *y*, respectively; finally, 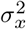 and 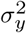 are the variances for the mutational distribution.

Intuitively, one may expect that increased environmental mismatch favors the evolution of cooperation, counteracting the decrease in population size. If there are rates of environmental change for which populations with a constant level of cooperation go extinct while those with an evolving level of cooperation do not, we would say that the evolution of cooperation facilitates evolutionary rescue. This process can indeed be observed for some life-cycles.

For example, Fig. 3A shows the evolutionary dynamics of a population belonging to a species with life-cycle D (Table 1). The pink curves depict the dynamics of a population where cooperation cannot evolve (or evolves much slower than the change in the environment). At the beginning of the numerical simulation, the population is perfectly adapted to the environment, and the level of cooperation is at equilibrium. As the optimum starts moving at a constant velocity, the population size decreases. For some time, adaptation in *y* keeps the population afloat, by lowering the rate of increase in the lag and the rate of decrease in population size. But as the population becomes ever smaller, adaptation decelerates, limited by the supply of beneficial m utations. The positive feedback between decreased population size and slower adaptation leads the population to extinction. Compare this with the blue curves, which depict the dynamics of a population where cooperation evolves relatively fast (parameters in the figure caption). The initial dynamics are very similar to the previous case: an increase in the lag and a corresponding decrease in population size. But this very decrease in population size rewards higher cooperative investments. As cooperation increases, it allows the population to grow and thus adapt faster to the environmental change. Confirming our intuition, the evolution of cooperation rescues the population and leads to a dynamical equilibrium (orange dot in Fig. 3A).

**Fig. 3.**
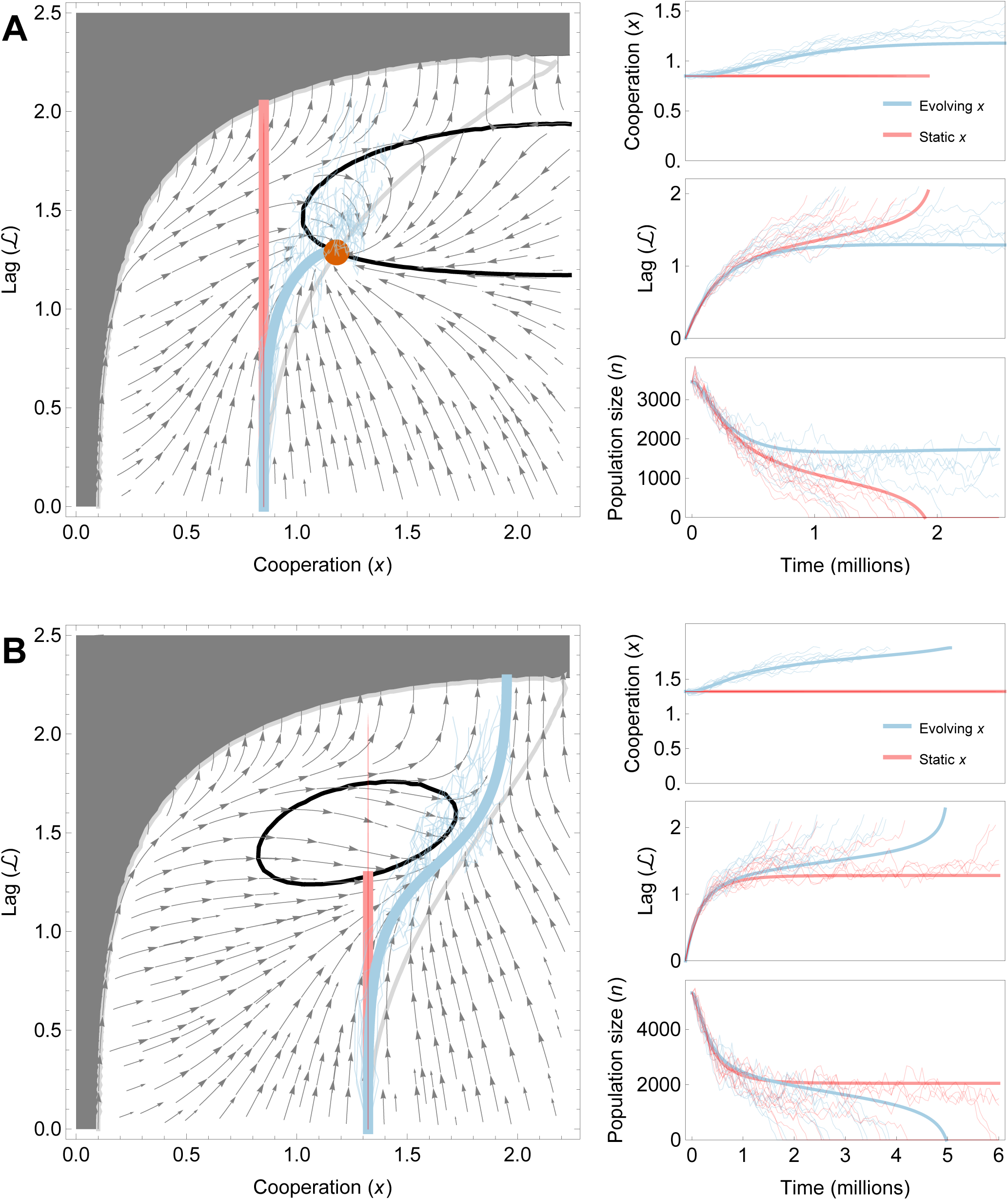
Evolution of cooperation with a moving optimum, for a species with life-cycle D (panel **A**) or life-cycle CD (panel **B**). The stream plot to the left indicates the change in a population’s lag 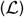 and cooperation (*x*) over time, for a population where cooperation evolves fast relative to the environmental change. The nullclines for lag 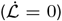 and cooperation (*ẋ*) are in black and light grey, respectively. Within the dark grey region, the population goes extinct. The thick blue curves are the result of numerical simulations, starting with zero lag and with *x* = *x*^⋆^. For comparison, the thick pink curves are the result of numerical simulations, with identical starting conditions, where cooperation does not evolve (*μ*_*x*_ = 0). The panels on the right show the time-dynamics of cooperation, lag, and population size for the same simulations. Thin lines show the mean cooperation value, mean lag, and population size of individual-based simulations (ten replicates for each treatment). Parameters: *r* = 10, *c* = 0.1, *s* = 1, *k* = 2, *μ*_*x*_ = *μ*_*y*_ = 10^−4^, *σ*_*x*_ = *σ*_*y*_ = 0.01; for **A**: *d* = 0.002, *v* = 5 × 10^−6^, *p* = 0.002; for **B:** *d* = 0.0019, *v* = 4.5 × 10^−5^, *p* = 5 × 10^−4^. In contrast to **A**, the parameters in **B** were chosen such that the nullclines do not intersect.

If the environment changes too fast, no dynamical equilibrium will exist. Higher environmental velocities result in a narrower lag nullcline (black line in Fig. 3A’s stream plot); if the velocity is too high, the nullcline vanishes, meaning that the population will fail to track the optimum and will go extinct regardless of the initial conditions (Fig. 4A). That said, species where cooperation can evolve will survive faster environmental changes compared to species where cooperation cannot evolve (Fig. 4A). Even when the velocities are low enough that both types of species can avoid extinction, species where cooperation can evolve will equilibrate at higher population sizes and lower lags (Fig. 4A).

**Fig. 4.**
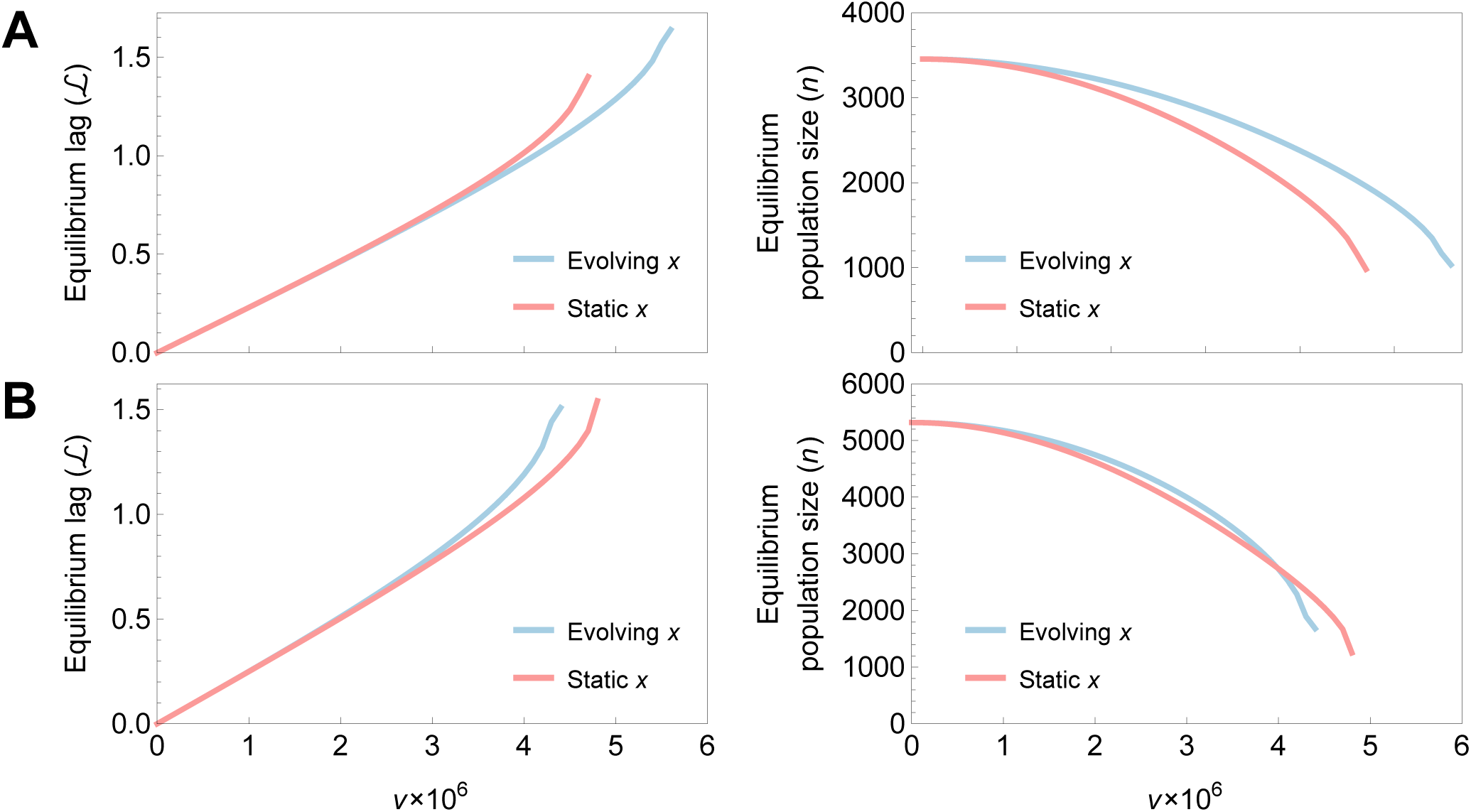
Lag (left) and population size (right) at the dynamical equilibrium, in populations with life-cycle D (panel **A**) or life-cycle CD (panel **B**), for different velocities of environmental change (*v*). The curves are interrupted at high velocities because the populations become extinct. Note that in **A**, for high values of *v*, the evolution of cooperation rescues populations that would otherwise undergo extinction; in contrast, in **B**, the opposite happens. Parameters: *r* = 10, *c* = 0.1, *s* = 1, *k* = 2, *μ*_*y*_ = 10 × 10^−4^, *σ*_*y*_ = 0.01; for **A**: *d* = 0.002, *p* = 0.002; for **B:** *d* = 0.0019, *p* = 5 × 10^−4^.

A dynamical equilibrium may also not exist if the game parameters do not favor the evolution of high cooperation. For example, larger interaction groups deter cooperation (because they decrease the fraction of investment benefits that return to the cooperator). Increasing *p* makes groups larger, and therefore it moves the cooperation nullcline (grey line in Fig. 3A’s stream plot) to the left, thus causing the dynamical equilibrium to vanish. Finally, even *if* a dynamical equilibrium exists, the population will be unable to reach it if it evolves too slowly in *x*. The pink curve in Fig. 3 represents the most extreme such scenario (*μ*_*x*_ = 0 or *σ*_*x*_ = 0). Note that the position of the equilibrium itself does not depend on *μ*_*x*_ or *σ*_*x*_, provided that *μ*_*x*_ > 0 and *σ*_*x*_ > 0.

Surprisingly, small differences in the life-cycle can result in a diametrically opposite outcome. Consider now a species with life-cycle CD (Table 1). Recall that the only difference between the two life-cycles is that in CD, the costs of cooperation decrease viability instead of fertility (as in D). In the one-dimensional dynamics, the differences between the two life-cycles are relatively minor (Fig. 2A): although adaptation is slightly slower for CD, the evolutionary equilibrium is the same in both life-cycles. However, the differences between the life cycles turn out to have important qualitative consequences for a population’s capacity to track a moving optimum.

In contrast to life-cycle D, the costs of cooperation decrease viability in CD, causing an increase in 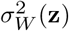 (Eq. 15), thus slowing evolution. The rate of evolution is proportional to 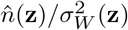; when costs do not affect viability, this quantity always increases with *x* < *x*_crit_ (purple curves in Table 1), but when they do, high values of cooperation cause slower rates of evolution. The changing environment causes the population to decline; cooperation becomes favored, but this increase in cooperation, while it may temporarily make the population larger (speeding adaptation by increasing the supply of mutations), also increases the variance in reproductive success, eventually making adaptation slower.

Qualitatively, this means that there are regions of parameter space where a population that would otherwise survive an environmental change—for example, a population whose mutation rate in *y* is high enough to keep up with the moving optimum, as in the pink curve in Fig. 3B—may *fail* to survive when cooperation evolves (blue curves in Fig. 3B). Counter-intuitively, the evolution of cooperation, which, for intermediate velocities, increases population size (relatively to the case where cooperation does not evolve, Fig. 4B), also dooms the population to extinction.

Thus, for some velocities of environmental change, the evolution of cooperation in CD leads to extinction (compared to populations where cooperation does not evolve, Figs. 3B and 4B). More generally, in stark contrast to D, the evolution of cooperation in CD results in higher equilibrium lags (compare Fig. 4B for CD to Fig. 4A for D). To understand the difference between D and CD, consider the change in lag, 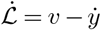. If a dynamical equilibrium exists, then at such an equilibrium, *ẏ* = *v*. From Eq. 15 we can write:

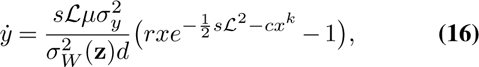

which, for life-cycled D and CD, equals:

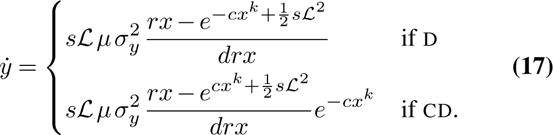

With life-cycle D, the maximum of *ẏ* occurs exactly at *x* = *x*_crit_. If there is a value of *x* for which *ẏ* = *v*, further increases in *x* will not compromise the population’s capacity to track the moving optimum, because *x* will never increase to values higher than *x*_crit_ (which maximizes population size). In contrast, for CD, *ẏ* is maximized at intermediate values of *x*, and then decreases. This means that, if evolution leads to a sufficiently high increase in *x*, the evolution of the functional trait will necessarily become slower than the velocity of environmental change. As cooperation slows adaptation in *y*, the population size declines, causing yet more cooperation to evolve. Thus, the evolution of cooperation promotes a vicious cycle that drives the population to extinction, an example of evolutionary suicide (Ferrière, 2000; Gyllenberg and Parvinen, 2001). For this reason, in contrast to D, the lag nullcline in CD (black line of Fig. 3B) vanishes for high values of cooperation. Of course, equilibria may still exist if the velocity is small, which widens the area enclosed by the lag nullcline, or if the game parameters are such that equilibrium cooperation is low. For example, for larger group sizes, *p*, cooperation is disfavored, so that the *x* nullcline in Fig. 3B moves to the left and may intersect the 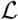 nullcline.

Eq. 16 has a maximum in 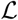, implying a saddle-node bifurcation in the dynamical equilibrium lag. This bifurcation corresponds to the fact that the equilibrium lag ceases to exist at high velocities (Fig. 4); it is an ‘evolutionary tipping point’ (Osmond and Klausmeier, 2017). This bifurcation also explains why simulated populations rapidly go extinct (Fig. 3). We can find the critical lag that maximizes the rate of evolution 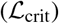 by calculating the root of 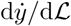:

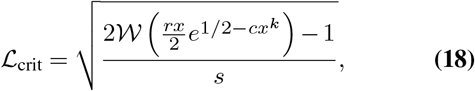

where 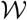 again denotes the Lambert *W*-function.

Having calculated the critical lag, we can plug it into Eq. 17 to obtain the maximum velocity of environmental change that can be tolerated by the population, before it goes extinct (for a given value of *x*). Although the resulting expression is not particularly insightful, a plot of this critical velocity confirms that increases in *x* < *x*_crit_ have qualitatively different effects for the life-cycles D and CD (Fig. 5).

**Fig. 5.**
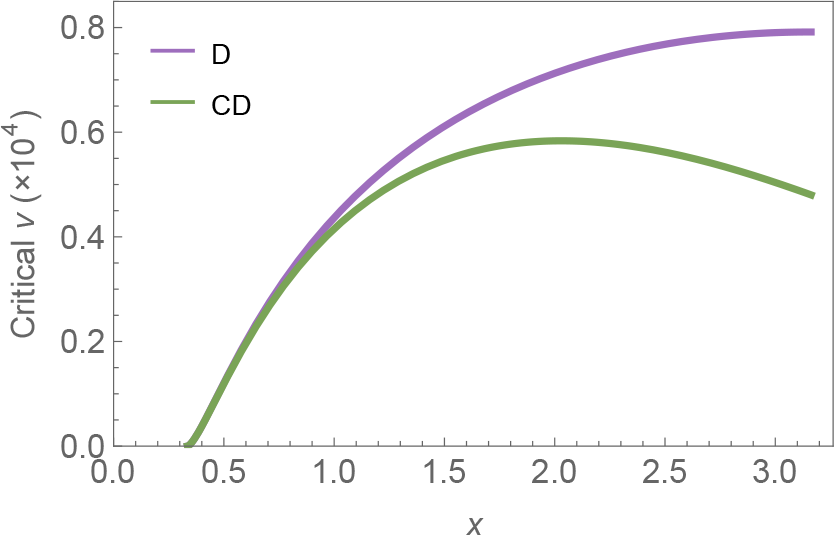
Critical velocity (maximal velocity of environmental change for which the population is able to track the moving optimum without undergoing extinction) as a function of the level of cooperative investment (*x* < *x*_crit_), for life-cycles D (purple) and CD (green). The evolution of increasing levels of cooperation makes populations able to endure faster rates of environmental change in D, but it has the opposite effect in CD. The critical velocity is calculated by subbing 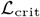 (Eq. 18) into *ẏ* (Eq. 17) Parameters: *d* = 0.015, *r* = 3, *c* = 0.05, *p* = 0.005, *k* = 2, *s* = 1.5, *μ*_*x*_ = *μ*_*y*_ = 5 × 10^−4^, *σ*_*x*_ = *σ*_*y*_ = 0.05.

In general, for any life-cycle where the costs of cooperation decrease viability, 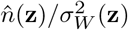 will decline as *x* approaches *x*_crit_, whereas for any life-cycle where the costs instead decrease fecundity, it will be maximized exactly at *x* = *x*_crit_. This is because the slope at 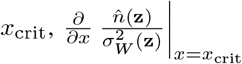, only depends on the derivative of 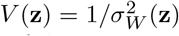 (since *n̂*(**z**) is maximized at *x*_crit_). The viability, in turn, can depend on *x* only through *C*(*x*) and/or *D*(*n̂*(**z**)). If the viability is affected by neither (life-cycle ∅), then its derivative will be zero. The same is true when the viability depends on *x* only through *D*(*n̂*(**z**)), again because *n̂*(**z**) is maximized at *x*_crit_. Thus, all life-cycles in which the costs of cooperation do not affect viability have a maximum at *x* = *x*_crit_. In contrast, when the viability is affected by the costs of cooperation, its derivative also depends on the derivative of (1 − *C*(*x*)), which is always negative. Therefore, if the costs of cooperation decrease viability, then 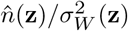 is declining with *x* at *x*_crit_, and increasing cooperation can slow evolution.

## 3. Discussion

In this manuscript, we have studied the role that the evolution of cooperation may play during adaptation to environmental change. By modeling cooperation as an ecological public goods game (Gokhale and Hauert, 2016; Hauert et al., 2006, 2008; Parvinen, 2010; Wakano et al., 2009), we were able to connect the dots between changes in environmental conditions and the evolution of social behavior. Our prediction was that changes in the environment would promote the evolution of cooperation, which would compensate for decreases in population size and permit populations to keep up with moving environments. When cooperation played out as a discrete game (as a result of the population branching into two strains, one consisting of cooperators and the other of defectors), we indeed observed these dynamics (see section S2 in the Supplemental Information). However, when cooperation is best described as a continuous game, we found out that widely different outcomes are possible, depending on the life-history of the species under study.

We contrasted a life-cycle where the costs of cooperation affect fecundity (D: Figs. 3A and 4A), to one where they affect viability (CD: Figs. 3B and 4B). In the former case the evolution of cooperation promoted rescue, while in the latter it led to evolutionary suicide. This seemingly paradoxical result arose because, in the latter case, while cooperation increased census population sizes, it also increased variance in reproductive success, thus declining the effective population size (Hedrick, 2005; Kimura and Crow, 1963) and slowing down adaptation by increasing the effect of genetic drift. The evolution of a population toward extinction—evolutionary suicide (Ferrière, 2000; Gyllenberg and Parvinen, 2001)—has often been discussed in the context of evolution favoring selfish individuals (Gyllenberg and Parvinen, 2001; Gyllenberg et al., 2002; Rankin et al., 2007), which leads to a tragedy of the commons (Hardin, 1968). However, here we have shown the opposite effect, where the evolution of increased levels of cooperation underlies evolutionary suicide.

Although we focused on models with non-overlapping generations, there are other possibilities. At the opposite end of the life-history spectrum, we may consider a species where generations overlap and in which individuals reproduce asynchronously by producing a single offspring at a time (as may be the case with microbes that reproduce by binary fission or by budding). Such a life-history resembles a continuous-time birth–death process, and it commonly modelled using the standard canonical equation of adaptive dynamics (Dieckmann and Law, 1996). A species with this life-history has a rate of evolution that is proportional to *n*/2 (Dieckmann and Law, 1996), rather than 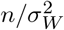, which is always increased by cooperation (up to *x*_crit_, which is the maximum value of cooperation that will evolve). Thus, such a species will qualitatively behave like a species with life-cycle D, meaning that the evolution of cooperation would favor persistence.

There is, of course, a wide range of possible life-histories between synchronous semelparity and asynchronous budding. Species may be iteroparous, having multiple broods throughout their lives. Furthermore, in iteroparous species reproduction may be either synchronous (as with many perennial plants) or asynchronous (as with most mammals). Iteroparous life-cycles may be more similar to CD-like models, because they incorporate reproductive variance. The canonical equation of adaptive dynamics for iteroparous diploids and haplodiploids (Metz and de Kovel, 2013) is similar to Eq. 13 in that it includes the term 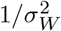, meaning that qualitatively, CD-type effects may be at play in such species (i.e., the evolution of cooperation may lead to evolutionary suicide in changing environments). The same is true for physiologically structured populations (Durinx et al., 2008).

Our results highlight that species engaging in cooperative interactions may exhibit disparate and counter-intuitive responses to environmental change. For example, mammals and birds engaging in public goods provisioning (although they are most often iteroparous) have a relatively low number of offspring, and low reproductive variance. In contrast, many social insects, which engage extensively in cooperative interactions, have non-overlapping generations and very high numbers of offspring, implying the possibility of high reproductive variance. Whether reproductive variance indeed increases with fecundity depends on the distribution of offspring number. In this model, we assumed that it does (by modeling fecundity as a Poisson distribution), which had a major influence in our results, but other choices could lead to different outcomes. Overall, whether the evolutionary dynamics of social behavior will ultimately be beneficial or detrimental during adaptation to environmental change is contingent on the species’ specific life history.

In the future, it would be interesting to consider whether these results apply also to other mechanisms of cooperation. For example, decreases in population size are associated with increased relatedness (Pepper, 2000), which could favor cooperation via kin selection. This provides the opportunity for dynamics very similar to the ones we have described. In contrast, other mechanisms of cooperation may affect adaptation in quite different ways. Particularly, in the case of between-species cooperation (mutualism), environmental change would set up coevolutionary feedback loops (Northfield and Ives, 2013) that are not well described by our framework.

## 4. Author contributions

GJBH conceived of the project, wrote the simulations, and wrote the paper. MMO supervised the project and contributed substantially to the simulations and to writing. Both authors participated in developing the model.

## 5. Acknowledgements

The authors are especially thankful to Michael Doebeli for thoughtful discussion and advice throughout the project’s development. We are also thankful to Leticia Avilés, Christoph Hauert, Koichi Ito, Ailene MacPherson, Sarah Otto, Linnéa Sandell, Sebastian Schreiber, and Michael Whitlock. R. Henriques created the bioR*χ*iv LATEXtemplate.

The authors acknowledge that this work was performed in the ancestral, unceded, and occupied territories of the Musqueam and Patwin peoples. The authors recognize Musqueam and Patwin land rights and sovereignty over their traditional homelands.

GJBH was funded by a Zoology Graduate Student Fellowship (Department of Zoology at UBC). MMO was funded by the National Science and Engineering Research Council (RGPIN862 2016-03711 to Sarah Otto), the National Institute of General Medical Sciences of the National Institutes of Health (NIH R01 GM108779 to Graham Coop), Banting (Canada), and the Center for Population Biology (University of California, Davis).

## 6. Data accessibility

Data and code, including a *Mathematica* notebook containing all calculations, are currently available in a directory hosted by the UBC Zoology department. The files will be moved to a permanent repository upon acceptance.

## Notes

https://cloud.zoology.ubc.ca/s/HfFFn8ASxYYMWJH

