## Supplemental Information for "During environmental change, cooperation can promote rescue or lead to evolutionary suicide"

September 17, 2019

#### S1 Canonical equation modified for non-overlapping generations

The canonical equation of adaptive dynamics, derived in Dieckmann and Law (1996), describes the dynamics of the mean evolutionary path that results from a continuous-time birth-death process. Here we follow the same steps to obtain a recurrence equation which resembles the canonical equation, but is more appropriate to describe populations with discrete, non-overlapping generations.

Let $x$ represent the current mean trait value of the evolving population (which is assumed to be quasi-monomorphic). The random variable $X(x', x)$ determines whether a newborn individual with trait value $x'$ will reach reproductive age. This will happen with probability $V(x', x)$, referred to as viability, so that $X(x', x) \sim \text{Bernoulli}(V(x', x))$. When she reproduces, her expected number of offspring (fertility) will be $F(x', x)$; the realized number of offspring is a random variate drawn from the random variable $Y(x', x)$. Following conventional practice, we assume in the main manuscript that $Y(x', x) \sim \text{Poisson}(F(x', x))$, but the appropriate distribution will depend on the species being studied (for example, in vertebrate species, the most appropriate distribution to describe variation in number of offspring may be the two-parameter Poisson-Consul distribution, also known as generalized Poisson, Kendall et al. 2010). Individuals are semelparous (they die upon giving birth). Thus, the Wrightian fitness is $W(x', x) = V(x', x) \cdot F(x', x)$.

#### S1.1 Stochastic description of trait substitution sequences

We assume that a successful mutation will rise to fixation before a new mutation occurs, such that no more than one mutation segregates in the population. In reality, even the most successful mutations will take many generations to achieve fixation, but we model the process as if successful mutations achieve fixation instantly. This corresponds to a separation of time-scales between ecological and evolutionary dynamics (Geritz et al. 1998).

Based on these assumptions, the evolutionary dynamics consists of a sequence of substitution events in which a mutant $x'$ replaces a resident $x$. Dieckmann and Law (1996) call this the *trait substitution sequence*. Because mutation and selection depend only on the present state of the population, the trait substitution sequence is Markovian. To describe this sequence of substitutions, we first need to calculate the probability per generation of the trait substitution $x \to x'$, which we call transition probability, $P_{x',x}$.

##### S1.1.1 Transition probabilities

Based on these assumptions, the transition probability per generation from $x \to x'$ is given by two factors: first, the probability per generation $\mathcal{M}(x', x)$ that the mutant enters the population; second, the probability of fixation $\mathcal{S}(x', x)$:

$$P_{x',x} = \mathcal{M}(x', x) \cdot \mathcal{S}(x', x). \tag{S1}$$

In a given generation, mutations can occur in any newborn individual. The number of newborns is $W(x, x)\hat{n}$, where $\hat{n} \equiv \hat{n}(x)$ is the equilibrium population size (censused before viability selection). The fraction of births that give rise to mutations is $\mu$. The probability distribution function of mutant trait values around the resident is $M(x' - x)$. Collecting these terms we obtain

$$\mathcal{M}(x', x) = W(x, x)\hat{n}\mu M(x' - x) \tag{S2}$$

for the probability per generation that the mutant enters the population.

The probability of fixation $\mathcal{S}(x', x)$ depends on demographic stochasticity. Mutants appear as single individuals, and thus fixation is dependent on avoiding stochastic extinction when rare. We make two assumptions: first, that the resident population size $\hat{n}$ is large enough that there's a negligible risk that the population will go accidentally extinct; second, that as long as an advantageous mutant escapes stochastic extinction, it will achieve fixation ('invasion implies fixation').

The probability of avoiding stochastic extinction can be calculated using discrete-time branching process theory (Allen 2010). The key parameters are the expectation of the per capita number of mutant offspring—i.e., the mutant's Wrightian fitness, $W(x', x)$—and the variance of the same quantity—i.e., the mutant's reproductive variance, $\sigma_W^2(x', x)$:

$$W(x', x) = \mathrm{E}[X(x', x)Y(x', x)] = V(x', x)F(x', x) \tag{S3}$$

$$\sigma_W^2(x', x) = \mathrm{Var}[X(x', x)Y(x', x)]. \tag{S4}$$

We will also represent the resident's reproductive variance as $\sigma_W^2 \equiv \sigma_W^2(x, x) = \mathrm{Var}[X(x, x)Y(x, x)]$.

The fixation probability, for a mutant that starts as a single copy, is (following Allen 2010):

$$\mathcal{S}(x', x) = \begin{cases} 1 - \exp\left(-\dfrac{2(W(x', x) - 1)}{\sigma_W^2(x', x)}\right) & \text{if } W(x', x) > 1 \\ 0 & \text{otherwise.} \end{cases} \tag{S5}$$

Eq. S5 is also used in other variants of the canonical equation of adaptive dynamics (e.g., Durinx et al. 2008; Metz and Kovel 2013).

###### S1.1.2 Stochastic dynamics

Using the transition probabilities, we can describe the dynamics of $x$ in evolutionary time. We define the transition rate $p_{x',x}$ as the limit of $P_{x',x}/\Delta t$ for small $\Delta t$. Given that the unit of time is one generation (i.e., $\Delta t = 1$), which is very small in relation to the evolutionary time scale, $p_{x',x} \approx P_{x',x}$, we can describe the dynamics of $x$ in continuous time as:

$$\frac{\mathrm{d}}{\mathrm{d}t} \Pr(x, t) = \int P_{x,x'} \Pr(x', t) - P_{x',x} \Pr(x, t)\, \mathrm{d}x', \tag{S6}$$

where $\Pr(x, t)$ denotes the probability that the trait value is $x$ at time $t$.

#### S1.2 Mean path

Imagine a large number $N$ of trait substitution sequences, $x_1(t), x_2(t), \ldots, x_N(t)$. The mean path is then defined as:

$$\langle x\rangle(t) = \int x \Pr(x,t)\mathrm{d}t. \tag{S7}$$

We can thus describe the dynamics of the mean path as:

$$\begin{aligned}\frac{\mathrm{d}}{\mathrm{d}t}\langle x\rangle &= \int x(\Pr(x,t+1) - \Pr(x,t))\mathrm{d}t \\ &= \int x \int P_{x,x'} \Pr(x',t) - P_{x',x}\Pr(x,t)\mathrm{d}x'\mathrm{d}x \\ &= \iint (x'-x)P_{x',x}\Pr(x,t)\mathrm{d}x'\mathrm{d}x.\end{aligned} \tag{S8}$$

Eq. S8 is, by definition, the mean of the quantity called the *first jump moment*,

$$a(x) = \int (x'-x)P_{x',x}\mathrm{d}x'. \tag{S9}$$

Thus,

$$\frac{\mathrm{d}}{\mathrm{d}t}\langle x\rangle = \langle a(x)\rangle(t). \tag{S10}$$

As long as the deviations of the stochastic path from the mean path are relatively small, the first jump moment will be an almost linear function of $x$, such that we can approximate the mean path as:

$$\frac{\mathrm{d}}{\mathrm{d}t}\langle x\rangle \approx a(\langle x\rangle(t)). \tag{S11}$$

We call this approximation the *deterministic path*.

#### S1.3 Deterministic path

Substituting Eq. S7 into Eq. S11, using Eqs. S1, S2, and S5 (and dropping the angle brackets) we obtain:

$$\frac{\mathrm{d}x}{\mathrm{d}t} = W(x,x)\hat{n}\mu \int_+ (x'-x)M(x'-x)\left[1 - \exp\left(-\frac{2(W(x',x)-1)}{\sigma_W^2(x',x)}\right)\right]\mathrm{d}x', \tag{S12}$$

where the range of integration has been restricted to advantageous mutations (i.e., $W(x',x) > 1$).

Note that the rate of evolution depends on the fertility and viability of all possible advantageous mutant trait values, $x'$, as is clear from the range of integration. In order to transform this global coupling into a local one we apply a Taylor expansion to the probability of fixation (the term within square brackets), when $x' \approx x$:

$$1 - \exp\left(-\frac{2(W(x',x)-1)}{\sigma_W^2(x',x)}\right) \approx (x'-x)\frac{2}{\sigma_W^2}\frac{dW(x',x)}{dx'}\bigg|_{x'=x}. \tag{S13}$$

This approximation implies that the mutational step size $(x'-x)$ is very small compared to the other terms. In particular, the approximation ceases to be valid if the reproductive variance $\sigma_W^2$ approaches the order of the mutation size.

Substituting this result into Eq. S12:

$$\frac{\mathrm{d}x}{\mathrm{d}t} = 2\frac{1}{\sigma_W^2}\hat{n}\mu\frac{\mathrm{d}W(x',x)}{\mathrm{d}x'}\bigg|_{x'=x}\int_+ (x'-x)^2 M(x'-x)\mathrm{d}x'. \tag{S14}$$

Finally, we focus on the mutation process. As long as $\mathrm{d}W/\mathrm{d}x$ is not zero there is directional selection to leading order; then, if $M(x'-x)$ decays fast enough as $x'$ departs from $x$ (i.e., small mutations), we only need to consider the effect of selection up to leading order, which means that, since mutations are symmetric about $x$, mutations are advantageous with a probability of $1/2$. Since the density of beneficial and deleterious mutations are equal, we can remove the restriction to the domain of integration by multiplying the right-hand side of Eq. S14 by $1/2$. By definition, the second moment of $M(x'-x)$ is the variance in mutation size, $\sigma_\mu^2 = \int (x'-x)^2 M(x'-x)\mathrm{d}x'$. This concludes our calculation:

$$\boxed{\frac{\mathrm{d}x}{\mathrm{d}t} = \frac{1}{\sigma_W^2}\hat{n}\mu\sigma_\mu^2\frac{\mathrm{d}W(x',x)}{\mathrm{d}x'}\bigg|_{x'=x}.} \tag{S15}$$

The main qualitative difference between this and the traditional (overlapping generations) canonical equation of adaptive dynamics (Dieckmann and Law 1996) is that the rate of evolution is inversely proportional to the reproductive variance (i.e., the variance of the per capita number of offspring). This is similar to other variants of the canonical equation of adaptive dynamics (e.g., Durinx et al. 2008; Metz and Kovel 2013).

If we assume that the number of offspring follows a Poisson distribution, $Y(x',x) \sim \text{Poisson}(F(x',x))$, then:

$$\sigma_W^2(x',x) = W(x',x)(1 + F(x',x) - W(x',x)). \tag{S16}$$

It follows that, at ecological equilibrium, $\sigma_W^2 = F(x,x)$. Then the rate of evolution is proportional to $1/F(x,x)$ (or equivalently, since $W(x,x) = V(x,x) \cdot F(x,x) = 1$, to the viability, $V(x,x)$).

### S2 Some remarks on post-branching (polymorphic) dynamics

As described in section 3.3.1 of the main text, the singular point $x^\star$ can be a branching point if it obeys Eq. 12 (main text). If that is the case, then after converging to the singular point, the population will branch into two separate strains (with trait values $x_1$ and $x_2$), which will evolve in opposite directions (Fig. 2B, main text). Individual-based simulations illustrate these two behaviors (Fig. S1).

As mentioned in the main text, individual-based simulations indicate that the parameter range under which evolutionary branching occurs and results in a stable dimorphic equilibrium is narrower than the analytical predictions suggest. In particular the population size of the cooperator is often so small that stochastic extinction of that branch quickly occurs. Nonetheless, for the sake of completeness, here we present some numerical results regarding the deterministic dynamics after branching.

#### S2.1 Polymorphic dynamics for a constant lag

For simplicity, we will first focus on the case of a constant lag, $\mathcal{L}(y) = 0$, in a constant environment, $v = 0$. Because we are concerned with adaptation in $x$ and the lag is fixed, we will drop the dependence on $y$ for the remainder of this section.

The benefits of cooperation now depend on both phenotypes. In a group with $g_1$ type 1 residents and $g_2 = g - g_1$ type 2 residents the benefit to the focal individual (with phenotype $x'$) is:

$$B_2(x'|x_1, x_2, g_1, g_2) = \frac{r}{g_1 + g_2 + 1} (x' + g_1 x_1 + g_2 x_2). \tag{S17}$$

The probability $\Pr(g|n)$ that the interaction group size is $g = g_1 + g_2$ is the same as Eq. 1 (main text), and the probability of $g_1$ of those being of type 1 is:

$$\Pr(g_1|n_1, n_2) = \binom{g}{g_1} \left(\frac{n_1}{n-1}\right)^{g_1} \left(\frac{n_2 - 1}{n-1}\right)^{g-g_1}, \tag{S18}$$

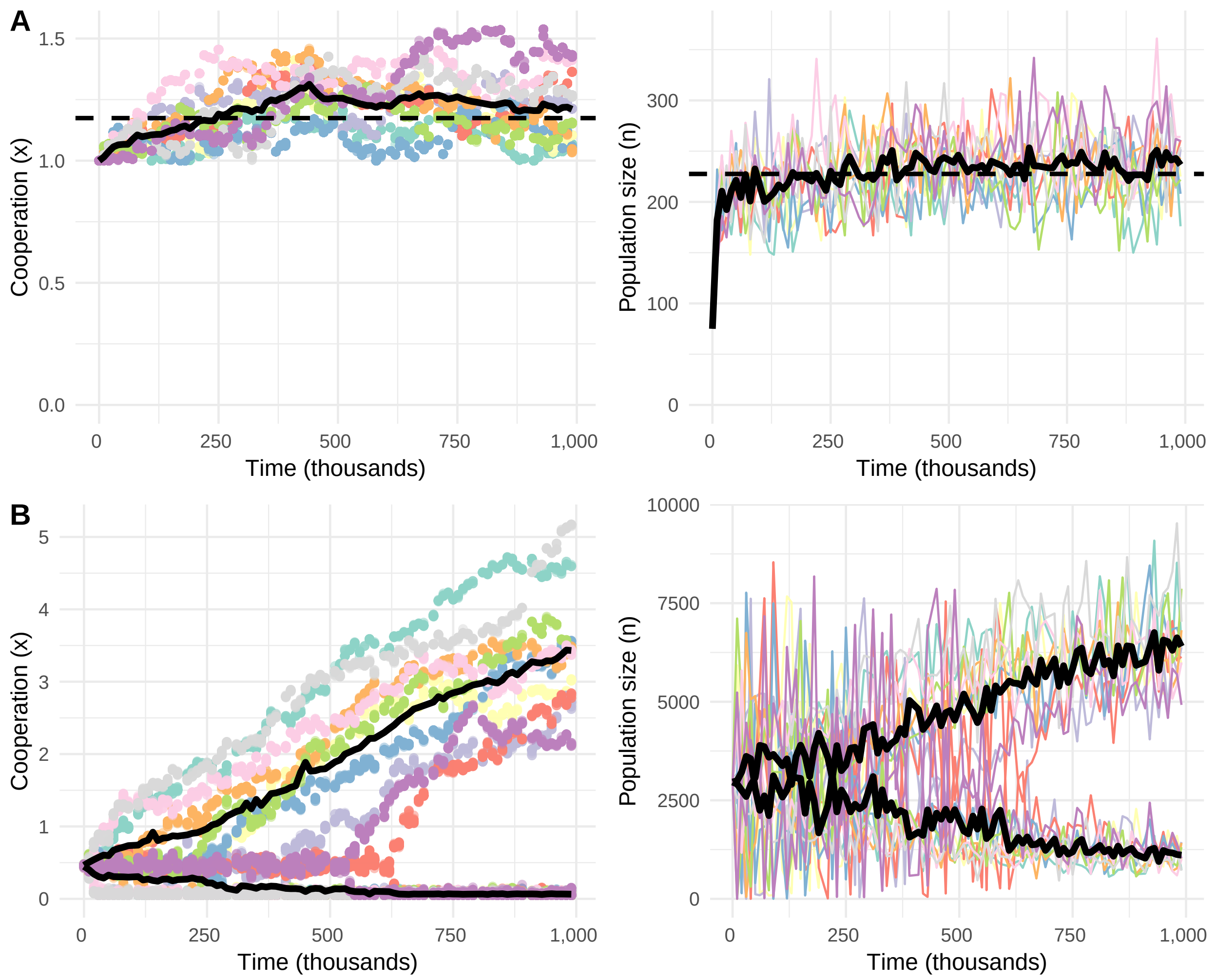


Figure S1: Individual-based simulations showing cooperation values (left) and population size (right) for both monomorphic and polymorphic dynamics, with static environments. **A:** Monomorphic approach to the equilibrium (indicated with a dashed line). **B:** Evolutionary branching, starting from the evolutionary branching point. In both panels, colors represent different replicates (10 replicates per panel). In the left-side panels, each dot indicates an individual (75 random individuals per replicate per time plot were plotted). Thick lines indicate strain averages; in **B**, individuals were assigned to different strains based on whether their trait value was higher or lower than the branching point. Parameters in **A**: $d = 0.01,\ r = 4,\ c = 0.3,\ k = 1.1,\ p = 0.01,\ \mu = 0.001,\ \sigma = 0.01,\ M = 1$; in **B**: $d = 5 \times 10^{-4},\ c = 0.05,\ r = 9,\ p = 0.01,\ k = 0.5,\ \mu = 0.01,\ \sigma = 0.01,\ M = 1$.

where $n_1$ and $n_2 = n - n_1$ are the population sizes for each strain. Therefore, the lifetime average benefit is:

$$\begin{aligned} \bar{B}_2(x'|x_1, x_2, n_1, n_2) &= \sum_{g=0}^{n-1} \sum_{g_1=0}^{g} \Pr(g|n)\ \Pr(g_1|n_1, n_2)\, B_2(x'|x_1, x_2, g_1, g_2) \\ &= \frac{r}{n(n-1)p}\left[x'(1-n)((1-p)^n - 1) + ((1-p)^n + np - 1)(n_1 x_1 + (n_2 - 1)x_2)\right]. \end{aligned} \tag{S19}$$

The lifetime average benefits allow us to calculate the Wrightian fitnesses in the dimorphic population, $W(x'|x_1,x_2) = \bar{B}_2(x'|x_1,x_2,\hat{n}_1,\hat{n}_2)(1-C(x'))D(\hat{n})M$, as well as the selection gradients for each of the two strains, $\mathcal{S}(x_i) = \partial W(x'|x_1,x_2)/\partial x'|_{x'=x_i}, i \in \{1,2\}$. We use these gradients to numerically evaluate the evolutionary dynamics of $x_1$ and $x_2$, using a coupled version of Eq. 13 (main text), see Fig. 2B.

After branching, the strategies diverge until the edge of the phenotype space is reached at $x_1^\star = 0$. In time, the other strain also reaches an equilibrium $x_2^\star$ (Fig. 2B, main text). At this point, the population will consist of a cooperator strain and a defector strain, which does not contribute to public good production. Thus, starting from a continuous phenotype space, we observe the evolutionary origin of defectors and cooperators (a tragedy of the commune, Doebeli et al. 2004).

The equilibrium value of cooperation $x_2^\star$ can be found by setting $x_1^\star = 0$. Then, the cooperator's selection gradient simplifies to:

$$\mathcal{S}(x_2) = \frac{1-C(x_2)}{x_2}\left(\frac{W(x_2|0,x_2)}{1-C(x_2)}(1-ckx_2^k) - W(0|0,x^2)\right). \tag{S20}$$

At equilibrium, $\mathcal{S}(x_2^\star) = 0$ and the fitnesses of both strains are one, so that the solution is determined by:

$$ck(x_2^\star)^k = C(x_2^\star), \tag{S21}$$

which depends only on $c$ and $k$. In particular, the equilibrium investment of the cooperator strain does not depend on the magnitude of that lag. Thus, when populations consist of two discrete phenotypes (defectors and cooperators), they adapt to mismatches between the functional trait $y$ and the environmental optimum only by changing the relative frequencies of each strain.

Because of this property, once evolutionary branching has occurred and bimorphic evolutionary equilibrium has been reached, our model becomes identical to a *discrete* ecological public goods game. The same model, then, permits us to investigate the interaction between environmental change and cooperation for both discrete and continuous types of cooperation.

For any given lag, at ecological equilibrium, $W(x_1|x_1,x_2) = W(x_2|x_1,x_2) = 1$, we can obtain a (numerical) solution for the strain sizes. The overall population size decreases with lag (Fig. S2B, blue curve), but the number of cooperators is higher at large lags (Fig. S2A). This change in the frequencies of cooperators and defectors allows populations to withstand much larger lags, and at higher population sizes, than they

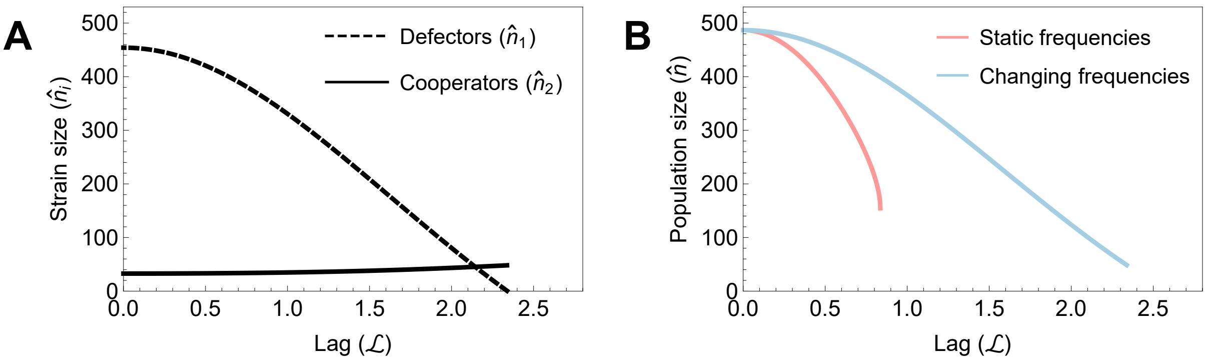


Figure S2: Equilibrium strain sizes in a bimorphic population with a constant lag. **A:** The equilibrium frequency of cooperators (solid line) is higher for large lags; however, since the frequency of defectors (dashed line) decreases, the total population size (blue line in **B**) is smaller at large lags. Nonetheless, evolution in the relative frequencies of cooperators and defectors allows populations to persist at much higher lags compared to a population where the frequency of cooperation is not an evolving trait (pink line in **B**). Parameters: $d = 0.01,\ r = 4,\ c = 0.3,\ k = 0.5,\ p = 0.01$.

would if cooperation simply occurred at a given (constant and lag-independent) frequency. The pink curve in Fig. S2B illustrates the population size for a reference population where the frequency of cooperators is kept constant at the same value that is displayed by the evolving population at equilibrium (blue curve) when $\mathcal{L} = 0$.

At very high lags, extinction occurs for the defector strain (Fig. S2A). When this happens, the population reverts back to the single strain case. Monomorphic dynamics will drive the population back to the evolutionary branching point, and branching will occur again, resulting in cycles of diversification and extinction.

#### S2.2 Polymorphic evolution during environmental change

We now turn to the evolution in the frequency of cooperators when the environment is changing at some velocity $v$, and populations consist of two discrete strains (cooperators and defectors).

We are interested in tracking the evolution in the functional trait $y$, whose selection gradient is given by Eq. 14 (main text). A dynamical equilibrium will exist whenever the lag becomes constant, i.e. $\dot{\mathcal{L}} \equiv v - \dot{y} = 0$. We only need to keep track of the sizes $\hat{n}_1, \hat{n}_2$ of each strain, since the trait values $x_1 = 0$ and $x_2 = x_2^\star$ (Eq. S21) do not evolve. However, we do need to keep track of each strain separately, because (since they have different sizes and payoffs) they evolve at different rates.

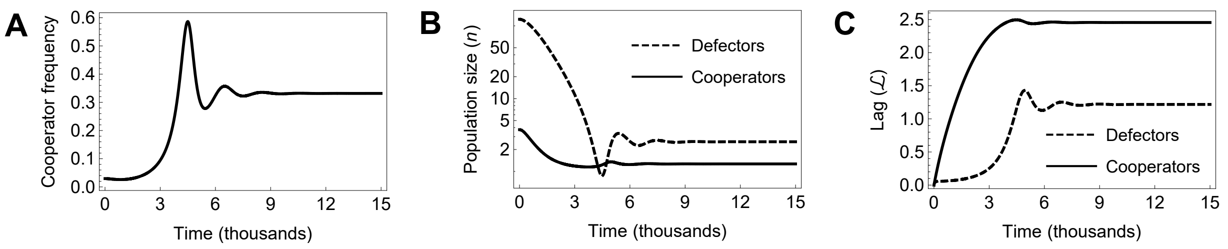


Figure S3: As the environment changes, the frequency of cooperators increases (**A**) which, after a period of oscillations, stabilizes the population size (**B**) and the distance to the optimum (**C**). Parameters: $d = 0.01,\ r = 4,\ c = 0.3,\ s = 1,\ p = 0.1,\ k = 0.5,\ \mu_y = 0.05,\ \sigma_y = 0.1,\ v = 0.0015$. Life-cycle: D.

The differential equations for the dynamics of the two strains' lags are given by:

$$\dot{\mathcal{L}}_i = v - \frac{1}{\sigma_W^2(y_i)}\hat{n}_i\mu_y\sigma_y^2 s\mathcal{L}_i,\ i \in \{1,2\} \tag{S22}$$

where strain sizes, $\hat{n}_i$, are the solution to $W(\mathbf{z}'_i|\mathbf{z}_1,\mathbf{z}_2) = \bar{B}_2(x'_i|\mathbf{z}_1,\mathbf{z}_2)(1-C(x'_i))D(\mathbf{z}_1,\mathbf{z}_2)M(y'_i) = 1,\ \forall i \in \{1,2\}$.

Numerically evaluating Eq. S22 shows that, as the environment changes, the distance to the optimum becomes much higher for cooperators than for defectors, which is not surprising since the cooperator strain size always seems to be smaller than the defector strain. Nonetheless, increasing distances from the optimum lower population size and hence favour cooperators, increasing their frequency. As an example, Fig. S3 shows both the population size and the lag becoming stabilized after a period of oscillatory behavior, achieving dynamical equilibrium (Fig. S3). The dramatic effects of life-cycle choice seen in the continuous games case are no longer present, because the value of $x$ is not evolving (nonetheless, life-cycle choice does affect the relative rate of evolution of the two strains).

Although the equilibrium cooperator frequency is much higher at fast velocities than at slow velocities, there can be a very slight decrease in the cooperator frequency at intermediate velocities of environmental change (Fig. S4A). This reflects the tension between opposing effects of environmental change. On the one hand, the decline of population size (Fig. S4B) favours cooperation; on the other hand, the smaller population size of the cooperators means a slower mutational input and larger lag (Fig. S4C). At either extreme (slow and fast velocities) cooperators will therefore equilibrate at higher relative frequencies when compared to intermediate velocities. At even higher velocities of environmental change, one of the strains goes extinct *before* any equilibrium is reached. In the numerical simulations we have attempted, this is always the defector

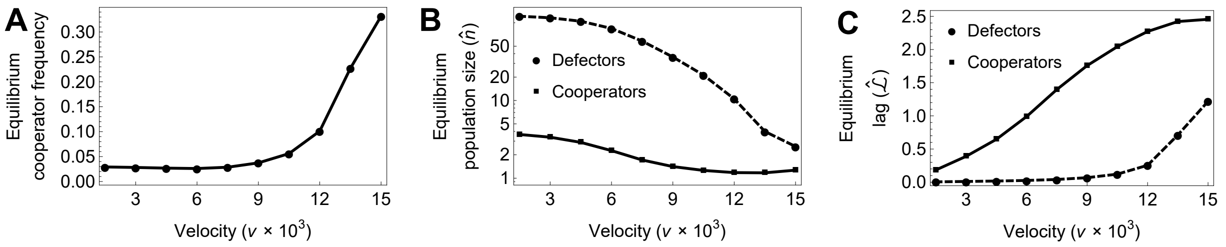


Figure S4: Frequency of cooperators (**A**), strain sizes (**B**) and distance to the optimum (**C**) at dynamical equilibrium, for different velocities of environmental change ($v$). Parameters: $d = 0.01,\ r = 4,\ c = 0.3,\ s = 1,\ p = 0.1,\ k = 0.5,\ \mu_y = 0.05,\ \sigma_y = 0.1$. Life-cycle: D. (In contrast to the continuous games case, the choice of life-cycle has no qualitative effect on this outcome, because the value of $x$ is not evolving.)

strain, because it experiences big oscillations in population size (as seen in Fig. S3B). After the extinction of the defector strain, the system reverts back to the monomorphic case, explored in detail in the main text. In our numerical results (such as the examples illustrated in Fig. S4) this reversal to monomorphism occurs at such high velocities of environmental change that no dynamical equilibrium can be reached, and the population goes extinct.
